# Comparison of Neural Tracking and Spectral Entropy in Patients with Disorders of Consciousness

**DOI:** 10.1101/2025.04.21.649509

**Authors:** Rien Sonck, Steven Laureys, Peter Diels, Tom Francart, Jonas Vanthornhout

## Abstract

**Objectives:** This study aims to explore the brain responses of patients with disorders of consciousness (DoC) to natural speech. Specifically, it focuses on a key characteristic of natural speech: the speech envelope. To achieve this, we employed two distinct measures. The first evaluates how effectively the patients’ brain activity “tracks” the speech envelope, called neural tracking. The second assesses the complexity of the brain’s responses to the speech stimulus, measured through spectral entropy. These two measures are then compared in their association with the patient’s clinical diagnosis and their level of behavioral responsiveness, both of which were assessed using the Coma Recovery Scale-Revised (CRS-R).

**Design:** Four patients with DoC participated in this study, during which their brain activity was recorded using electroencephalography (EEG). At the same time, they listened to a narrated story in both Dutch and Swedish. Additionally, EEG baseline recordings were collected. We employed a backward modeling approach to evaluate the speech envelope’s neural tracking. This technique involves training a model to map the relationship between EEG signals and the corresponding speech envelope. Once the model is trained, it can use unseen EEG data to reconstruct the speech envelope, which is then compared to the original speech envelope to assess how effectively the patient processed the auditory stimulus. For the behavioral assessment, we recalculated the CRS-R score of each patient into the CSR-R index, a more meaningful score that utilizes all the information contained in the CRS-R instead of only the highest scores on each subscale.

**Results:** Our findings revealed positive correlations between spectral entropy and the CRS-R index, which were more pronounced during the listening conditions than the baseline. While neural tracking of the speech envelope did not correlate with the CRS-R index, it did exhibit a positive association with CRS-R diagnoses, indicating that patients with better clinical diagnoses demonstrated higher levels of neural tracking. Additionally, we identified an interaction effect between spectral entropy and neural tracking. Specifically, higher levels of neural tracking were associated with a stronger positive relationship between spectral entropy and the CRS-R index. In contrast, when neural tracking was lower, this relationship disappeared.

**Conclusion:** This study demonstrated the potential of neural tracking and spectral entropy as complementary tools to investigate patients with DoC. Spectral entropy proved valuable for assessing behavioral responsiveness, while neural tracking shows promise in assessing the DoC diagnosis. **Terms**: disorders of consciousness (DoC), neural tracking, speech envelope, spectral entropy

## 1. Introduction

After an injury or acute brain disease, patients can enter a state of diminished or absent awareness, broadly classified as a disorder of consciousness (DoC). The accurate diagnosis of DoC is challenging, as it requires differentiating between levels of awareness. Patients with no signs of awareness are diagnosed with nonresponsive wakefulness syndrome (UWS). Patients with UWS are characterized by spontaneous eye opening and the presence of a sleep-wake cycle, but without evidence of intentional interaction with their environment. In contrast, patients with limited but reproducible awareness are classified as being in a minimally conscious state (MCS), where they may exhibit inconsistent behaviors such as tracking an object or responding to certain commands. However, they remain unable to communicate effectively (Giacino et al., 2002; Laureys et al., 2010).

Misdiagnosis between these states is alarmingly common, with studies reporting error rates as high as 33–43% (Schnakers et al., 2009; Stender et al., 2014). However, tools such as the Coma Recovery Scale-Revised (CRS-R) have improved diagnostic precision. The CRS-R assesses responses to stimuli via behaviors such as visually tracking an examiner’s hand or responding to tactile commands. Crucially, ‘command following’ - the ability to execute simple commands - is a clinical milestone that distinguishes MCS from UWS, with early evidence of command-following related to better recovery outcomes (Giacino et al., 2004; Whyte et al., 2009). However, a key limitation remains: the absence of observable behavioral responses does not necessarily imply an absence of awareness. Some patients retain covert cognitive capacities that are undetectable through behavioral measures (Laureys & Schiff, 2012; Sanders et al., 2012; Stender et al., 2014). This limitation led the field of DoC to explore and investigate neuroimaging tools as an objective motor-independent diagnostic tool. The main focus came on detecting ‘command following’ to be able to distinguish MCS from UWS (e.g., Cruse et al., 2012; Goldfine et al., 2011; Lulé et al., 2013; Monti et al., 2010; Owen et al., 2006; Pokorny et al., 2013; Schnakers et al., 2008).

However, failure to follow commands with neuroimaging does not necessarily indicate an absence of awareness and communication ability (Bodien & Giacino, 2016; Comte et al., 2015; Rashid et al., 2020; Stender et al., 2014) The success depends on patients actively engaging in tasks (Kleih et al., 2010; Nijboer et al., 2010), this faces challenges as patients with DoC are known to have fluctuating vigilance (Piarulli et al., 2016; Wannez et al., 2017) and may not always have the motivation required to maintain performance. Even among patients who do not follow instructions, there are significant differences in their levels of awareness and cognitive function. For example, patients with MCS are further categorized into MCS+ (with preserved language processing) and MCS-(without preserved language processing). MCS-patients exhibit greater functional impairments and lack command-following but may still respond to sensory input, such as tracking their reflection in a mirror (Thibaut et al., 2020). Some patients with UWS exhibit brain activity patterns similar to those of patients with MCS, despite the absence of overt behavioral responses. This subset of patients is classified as having cognitive motor dissociation (CMD), also called MCS* (Owen et al., 2006; Schiff, 2015; Thibaut et al., 2021). An accurate diagnosis of these subgroups is essential. Clearer distinctions between these groups could lead to tailored rehabilitation strategies, improved quality of life, and more reliable prognoses. Due to these subgroups, researchers are increasingly focused on assessing residual language processing to uncover covert cognitive abilities in non-communicating patients (Aubinet et al., 2022). Neural signatures of language processing provide a potential insight into the cognitive capacities of patients who appear to be behaviorally unresponsive, guiding rehabilitation strategies.

Paradigms that evaluate the brain’s capacity to track linguistic structures, such as hierarchical sentence processing and speech tracking, have shown promising results (Ding et al., 2016; Gui et al., 2020). These approaches go deeper than simple auditory-evoked responses by examining how the brain encodes and processes naturalistic speech. Previous research has primarily explored neural entrainment through coherence measures and frequency analysis (Jia et al., 2023; Xu et al., 2021).

Although previous studies have advanced our understanding of rhythmic alignment between brain activity and auditory stimuli, they have yet to fully address how the brain models the tracking of sound and speech. To fill this gap, the present study employs a backward-decoding model approach. This method offers a direct mapping between neural activity and speech features, enabling the reconstruction of stimulus features directly from brain responses. By quantifying how effectively the brain “tracks” specific speech features, this approach provides a powerful tool to investigate the neural mechanisms underlying continuous speech processing (Gillis et al., 2022).

In this study, we focus on the speech envelope, an ecologically speech feature that has already been extensively studied (e.g., Biesmans et al., 2017; Ding et al., 2014, 2016; Shannon et al., 1995), and also recently in DoC (Jia et al., 2023). The speech envelope are the low-frequency modulations in speech, which elicit corresponding low-frequency cortical responses in the auditory cortex. These neural responses reflect both bottom-up sensory processes and top-down processes related to speech intelligibility (Ding et al., 2014). Although the speech envelope is necessary for intelligibility, it is not sufficient on its own (Decruy et al., 2019; Kösem et al., 2023; Vanthornhout et al., 2018, 2019; Verschueren et al., 2022).

The goal of this study is twofold: first, to evaluate whether this modeling approach shows a relationship with diagnosis and behavioral responsiveness in patients with DoC, and second, to compare its performance with spectral entropy, a well-established measure of brain signal complexity. Spectral entropy has been widely used to assess the depth of anesthesia and to distinguish between conscious and unconscious states (Rezek & Roberts, 1998; Vakkuri et al., 2004; Viertiö-Oja et al., 2004). In the context of DoC, spectral entropy has shown promise as a tool for differentiating UWS from MCS. Patients with MCS exhibit higher spectral entropy values compared to those with UWS (Sitt et al., 2014), and spectral entropy has been found to positively correlate with the CRS-R score (Gosseries et al., 2011). Furthermore, it can detect arousal fluctuations within individual patients (Piarulli et al., 2016). Given its ability to differentiate UWS from MCS, its correlation with CRS-R scores, and its sensitivity to arousal changes, spectral entropy serves as a valuable benchmark to evaluate the performance of neural tracking.

## 2. Materials and methods

### 2.1. Patients

Five patients participated in this experiment. One patient was excluded due to poor electroencephalogram (EEG) quality, which was attributed to high interference that rendered neural responses unmeasurable. The age and sex of all patients are given in Table 1. Inclusion criteria required participants to be 18 years or older, at least 28 days post-injury, and to have a prior diagnosis of a DoC. Patients with a history of brain injury, hearing loss, or acute illness were excluded. Neuromuscular blocking agents or sedatives were not allowed within 24 hours prior to testing. Informed consent was obtained from a legal representative on behalf of each patient, with the approval of the Ethics Committee of the University Hospital of Liège (2020-173) and the Ethics Committee Research UZ/ KU Leuven (S64320). All patients are fluent in the Dutch language. Four patients were recruited and tested at the Inkendaal Rehabilitation Center, while the other patient was recruited and tested at the General Hospital Klina. Injury information for each patient is provided below.

**Table 1:** Patient information including demographics (a), diagnosis, and injury details (b). “Time since injury” refers to the duration between the injury and the first electroencephalography session. Abbreviation: MCS-, Minimally Conscious State minus; UWS, Unresponsive Wakefulness Syndrome; TBI, Traumatic Brain Injury; SCA, Sudden Cardiac Arrest.

| Subject | Sex | Age at time of testing |
| --- | --- | --- |
| Patient-01 | M | 21 |
| Patient-02 | M | 62 |
| Patient-03 | F | 57 |
| Patient-04 | M | 72 |

(a) Demographic information for each participant.
| Subject | Diagnosis | Time since injury | Injury |
| --- | --- | --- | --- |
| Patient-01 | MCS- | 15 weeks | TBI |
| Patient-02 | UWS | 33 weeks | SCA |
| Patient-03 | MCS- | 17 weeks | TBI |
| Patient-04 | MCS+ | 38 weeks | TBI |

(b) Injury and diagnosis details for each patient.

**Table 2:**
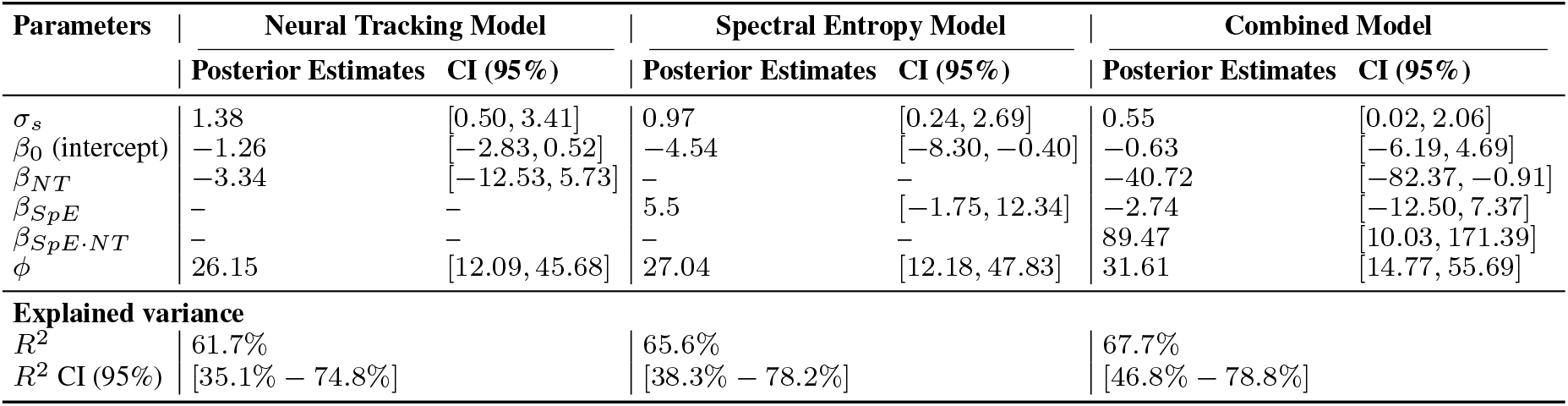
Results from the spectral entropy model (Eq. 4), the neural tracking model (Eq. 5), and the combined model, including both spectral entropy and neural tracking (Eq. 6). The table shows the posterior estimates and 95% credible intervals(CI) for each parameter across the three models. The explained variance section includes Bayes R^2^ and its 95% CI.

All patient injuries occurred in 2023 and early 2024. The exact dates are not given due to patient privacy.

#### 2.1.1. Patient-01: injury details

The patient was admitted to the hospital with craniocerebral trauma. Initial CT scans revealed limited subarachnoid bleeding in the right parietal region and minor intraventricular bleeding. Subsequent magnetic resonance imaging (MRI) indicated signs of diffuse axonal injury, with lesions identified in the centrum semiovale, splenium of the corpus callosum, and mesencephalon, as well as multiple contusions in the mesencephalon, bilateral basal forebrain, and left temporal lobe. A visual evoked potential test using flash stimulation showed bilateral responses, although with abnormal morphology in the left occipital region. Brainstem auditory evoked potentials demonstrated a normal brainstem response but revealed a delayed first peak when stimulating the left side, suggesting peripheral hearing impairment on that side. However, since the stimulation protocol used bilateral input, this impairment is expected to have a negligible effect on overall results. The patient was enrolled in the study at week 15, week 16, and week 17 post-injury.

#### 2.1.2. Patient-02: injury details

The patient was found to have facial trauma, along with a cardiorespiratory arrest of unknown duration, requiring prolonged resuscitation. As a result, the patient suffered brain damage due to oxygen deprivation (post hypoxic encephalopathy), which evolved into unresponsive wakefulness syndrome. CT scans did not show any signs of direct traumatic brain injury, but signs of hypoxic brain damage with temporary cerebral edema, hemorrhage in the nucleus lentiformis on the right side with evolution to hypodensities of the nucleus lentiformis on both sides. The EEG showed burst suppression patterns and spikes. Levetiracetam was started for clonic facial spasms. The patient was enrolled in the study at week 33, week 34, and week 36 postinjury.

#### 2.1.3. Patient-03: injury details

The patient was admitted to the hospital with severe polytrauma, including craniocerebral trauma. The injuries consisted of a subarachnoid hemorrhage in the right occipital lobe and a subdural hematoma-hygroma over the right convexity, accompanied by a mass effect and a midline shift to the left. To address these conditions, a temporary bilateral craniectomy was performed. An MRI scan revealed evidence of diffuse axonal injury and post-traumatic contusions on the right side, affecting the frontal, parietal, temporal, and occipital lobes, as well as the left frontal lobe. The patient was enrolled in the study at week 17, week 18, and week 25 post-injury.

#### 2.1.4. Patient-04: injury details

The patient, with a previous diagnosis of Parkinson’s disease, was admitted to the hospital due to neurotrauma. CT imaging revealed bilateral temporal subarachnoid hemorrhages, a left frontal intraparenchymal lesion, and a right anterolateral temporal lesion that could involve an epidural component. To address the epidural hematoma, the patient underwent a craniotomy to remove it. The patient was enrolled in the study at week 38, week 39, and week 44 post-injury.

### 2.2. Experimental procedure

The audio stimuli were presented binaurally using an RME Fireface UC soundcard (Haimhausen, Bayern, Germany) via ER-3A insert earphones (Etymotic Research Inc, IL, USA). Patients listened to the story “The Survivors” by Alex Schulman, presented in Dutch “De overlevenden” (20 minutes) or Swedish “ö verlevarna” (12 minutes), both narrated by the same female speaker. The audio of these stories was presented at 65 dBA using the APEX software (Francart et al., 2008).. Patients were tested in clinical facilities using a mobile electroencephalography setup, which allowed the test to be carried out in the patient’s room or any other room available in the facility. Each patient underwent three testing sessions, with at least one week between each session. For Patient-01, the data from the third session were removed because the cap became loose during the experiment. Each session began with a 5-minute baseline EEG recording, followed by a CRS-R assessment. For Patient-01, only one baseline was recorded per session. In contrast, for Patient-02, Patient-03, and Patient-04, baselines were recorded between each audio condition (see Figure 1). All patients were subjected to multiple audio conditions, although only the narrated story conditions are discussed here. In total, there were four narrated conditions: two Dutch conditions, each lasting 10 minutes, and two Swedish conditions, each 6 minutes long. Dutch and Swedish conditions alternated.

**Figure 1.**
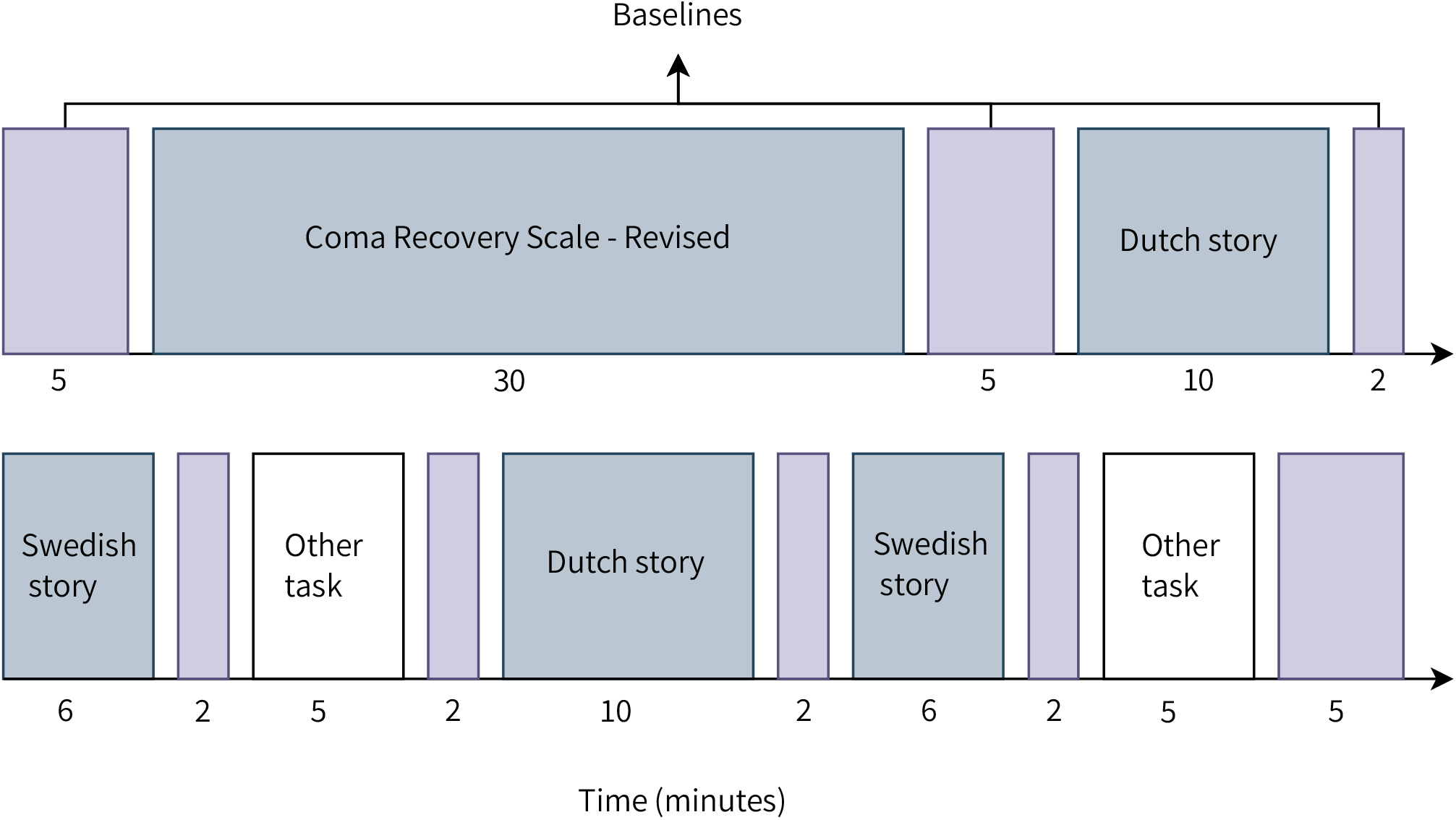
Timeline of one experimental session. Baseline EEG recordings were taken before and after each behavioral and EEG task.

### 2.3. EEG acquisition

We recorded the brain activity of each patient using electroencephalography at an 8192 Hz sampling rate with a 64-channel BioSemi ActiveTwo system and ActiView software (BioSemi, Amsterdam, The Netherlands). Ag / AgCl electrodes were placed on the head according to the 10-20 system (Oostenveld & Praamstra, 2001). Triggers were added to synchronize the audio stimuli with the EEG recording.

### 2.4. Signal preprocessing

EEG data preprocessing and analysis were performed offline using MATLAB (2023b). Data were filtered using a second-order zero-phase Butterworth high-pass filter in the backward and forward direction with a 0.5 Hz cutoff. Next, the data were downsampled to 256 Hz using the downsample function of MATLAB, which includes an anti-aliasing filter. A comb-notch filter was applied at 50 Hz and 100 Hz. Following these initial steps, preprocessing diverged according to the specific methods used: spectral entropy and neural tracking.

### 2.5. Outcome measures

Each electroencephalography session yields one CRS-R measurement, along with four measurements each for spectral entropy and neural tracking: two for Dutch listening and two for Swedish listening.

#### 2.5.1. Spectral entropy

Spectral entropy (SpE) quantifies the level of order or disorder in the frequency domain of a signal. SpE values range from 0 to 1, with values near 1 indicating a highly disordered or unpredictable signal, and values closer to 0 reflecting greater order or predictability.

##### Signal preprocessing

Bad channels identified during the EEG session were removed, and the remaining good channels were cleaned using Independent Component Analysis (ICA) using the infomax ICA algorithm (Bell & Sejnowski, 1995) in EEGLAB (Delorme & Makeig, 2004). This process resulted in 64 artifact components, minus the number of bad channels. These components were initially automatically classified using the ICLabel software (Pion-Tonachini et al., 2019), followed by manual inspection for verification. The non-brain components were discarded, and the brain components were projected back to channel space. Bad channels were then re-added and linearly interpolated. Finally, the data were re-referenced to Cz and downsampled to 128 Hz.

##### Calculation

To compute SpE, the time-frequency power spectrum for each channel was obtained using a Morlet wavelet transform for 0.5–32 Hz with 0.5 Hz increments, 7 cycles and 1-second nonoverlapping epochs. SpE was then calculated for each channel and epoch using the following formula (Viertiö-Oja et al., 2004):

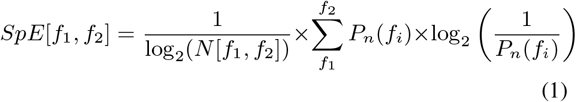

where

- [*f*_1_, *f*_2_] is the frequency range and *N* [*f*_1_, *f*_2_] is the total number of frequency components within this range.
- *f*_*i*_ represents each frequency component in the range [*f*_1_, *f*_2_].
- *P*_*n*_(*f*_*i*_) is the normalized power spectrum at frequency *f*_*i*_, calculated as the power at each frequency divided by the total power across all frequencies in the range [*f*_1_, *f*_2_].

#### 2.5.2. Neural tracking of the speech envelope

The primary objective is to explore the relationship between the EEG signals and the auditory stimulus presented during the recording. This can be achieved in two ways: by reconstructing the stimulus envelope from the EEG signals (backward/decoder model) or by reconstructing the EEG signals from the stimulus envelope (forward/encoder model). These approaches aim to determine how well neural activity “tracks” or aligns with the auditory stimulus (for a comprehensive overview, see Gillis et al., 2022). Here, we will focus solely on using a decoder model.

##### Signal preprocessing

Eye artifacts were removed using a multi-channel Wiener filter (Somers et al., 2018). Subsequently, the bad channels identified during the EEG session were linearly interpolated, and the data were re-referenced to the common average. The stimulus envelope was extracted using the method described by Biesmans et al., 2017, employing a gammatone filter bank and power-law compression. The same parameters as Vanthornhout et al., 2018 were used for the filter bank configuration. The envelope and corresponding EEG data were separately z-score normalized for each EEG condition. The data were then filtered using a high-pass filter with a cutoff frequency of 0.5 Hz and a low-pass filter with a cutoff frequency of 32 Hz. For each filter, a least squares filter was applied for each band, with a filter order of 2000. The filter design used stop-band attenuation of 80 dB and pass-band ripple of 1 dB. Finally, both the envelope and the EEG data were downsampled to 128 Hz.

##### Calculation

The decoder model *g* is trained with leave-one-out cross-validation, using 30-second folds of neural data to learn a set of weights that map the neural data to the stimulus envelope. The decoder model achieves this by minimizing the mean square error between the actual stimulus envelope *s*(*t*) and the reconstructed envelope *ŝ*(*t*) over time *t*:

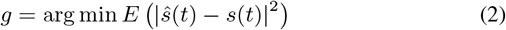

In practice, the decoder model achieves this by using ridge regression to solve for optimal weights (Machens et al., 2004). We used the same configuration as explained by Vanthornhout et al., 2018 with a post-stimulus integration window spanning 0 to 250 ms.

Once trained, the model can be applied to unseen neural data and reconstruct the corresponding stimulus envelope. The reconstructed stimulus is compared to the actual stimulus envelope using a Spearman correlation. The resulting correlation coefficient serves as the neural tracking value, with a higher correlation indicating that the model was more successful in reconstructing the stimulus envelope from the neural data, which can be seen as a marker of how intelligible the speech was to the participant (Vanthornhout et al., 2018).

#### 2.5.3. CRS-R index

The CRS-R is made up of six subscales: auditory function, visual function, motor function, oromotor function, communication, and arousal. Each subscale is scored based on the best behavior observed during the assessment. The total CRS-R score is obtained by summing the scores of all subscales. However, the total CRS-R score alone cannot be used to diagnose patients. The diagnosis depends on the specific types of behavior observed rather than the overall score. For example, two patients may both have a CRS-R score of 8, but one could be diagnosed with UWS, while the other could be diagnosed with MCS-. This distinction arises because behaviors like visual pursuit –a key diagnostic marker for MCS-– may be present in one patient but absent in the other.

To address the previous limitation, researchers have developed the CRS-R index to provide a more interpretable and diagnostically meaningful score (Annen et al., 2019). Unlike the total CRS-R score, which only considers the highest scores of each subscale, the CRS-R index incorporates all observed behaviors within each subscale. It assigns weights to these behaviors based on their diagnostic significance and normalizes scores across subscales to account for differences in maximum possible scores (e.g., auditory function has a maximum score of 4, while visual function has a maximum score of 5).

The CRS-R index is calculated by summing the weighted scores for all observed behaviors across subscales and transforming this sum into a scale ranging from 0 to 100. A cutoff score of 8.315 has been shown to reliably distinguish between patients with UWS and MCS (Annen et al., 2019).

### 2.6. Statistics

To investigate the relationship between spectral entropy, neural tracking of the speech envelope, CRS-R index outcomes, and CRS-R diagnoses, we employed two complementary approaches: Spearman correlations and Bayesian regression modeling.

Spearman correlations provided a rapid and straightforward assessment of the monotonic relationship between predictors and the outcome without assuming linearity. For visualization purposes, we plot this relationship using a second-degree polynomial. In contrast, Bayesian regression modeling offered a more robust framework by quantifying uncertainty in the estimated effects of spectral entropy and neural tracking on the CRS-R index and CRS-R diagnoses.

To ensure valid comparisons across conditions, we trimmed recording durations for the correlation analyses. For SpE calculations, both story listening conditions were trimmed to match the length of the baseline recordings (5-minutes). For neural tracking analyses, the Dutch listening recordings were trimmed to 6 minutes to match the duration of the Swedish listening recordings.

#### 2.6.1. Bayesian regression modeling: Spectral entropy versus neural tracking

We constructed several Bayesian regression models (BRM) using R (R Core Team, 2020) and the brms package (Bürkner, 2021) using only the 10-minute Dutch listening data. In each model, the dependent variables were the CRS-R index or the CRS-R diagnoses, and the predictors (fixed-effects) were spectral entropy and/or neural tracking. Our goal was to investigate how predictors relate to the CRS-R outcomes.

##### CRS-R Index model

The observed CRS-R indices in this study are skewed, strictly positive, and restricted between 0 and We scaled them to lie between 0 and 1. This scaling enables us to model the data appropriately using a Beta distribution:

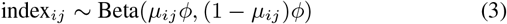

where:

- *index*_*ij*_ is the *i*-th observation of the CRS-R index of the *j*-th subject.
- *μ*_*ij*_ is the *i*-th predicted mean of the Beta distribution of the *j*-th subject..
- *ϕ* is the precision parameter of the Beta distribution, controlling the concentration of values around *μ*_*ij*_.

The mean parameter *μ*_*ij*_ is predicted by Bayesian regression models using a logit function:

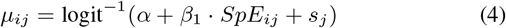

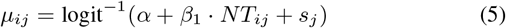

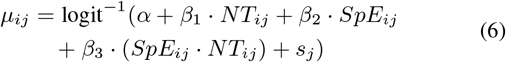

where:

- *β* are the regression coefficients.
- *NT* is the fixed effect for neural tracking.
- *SpE* is the fixed effect for the spectral entropy.
- *α* is the intercept.
- 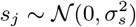 is the random intercept for subject *j*.

##### CRS-R diagnosis model

We model diagnosis as an ordinal outcome (UWS *<* MCS-*<* MCS+), using a Bayesian cumulative logit regression model:

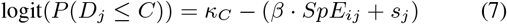

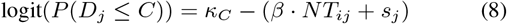

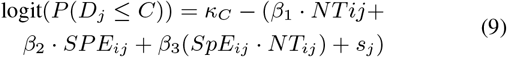

where:

- *D*_*j*_ is the best obtained diagnosis for subject *j*.
- *C* ∈*UWS, MCS*− represents the ordinal categories (with MCS+ as the reference level).
- *P* (*D*_*j*_ ≤ *C*) is the cumulative probability that subject *j*’s diagnosis falls in category *C* or lower.
- *β* are the regression coefficients.
- *NT* is the fixed effect for neural tracking.
- *SpE* is the fixed effect for the spectral entropy.
- *κ*_*C*_ are the threshold parameters (intercepts) for each cumulative probability boundary.
- 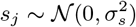 is the random intercept for subject *j*. Due to convergence issues, the CRS-R diagnosis model incorporating interaction effects, Equation 9, was excluded from further analyses. A detailed description of the Bayesian models and their specifications can be found in Appendix A.

## 3. Results

### 3.1. Behavioral results for each patient

Figure 2 summarizes the CRS-R sessions conducted by both the nursing staff in the rehabilitation centers and our team. Each patient is accompanied by a brief summary based on observations from CRS-R assessments and clinical neurological examinations.

**Figure 2.**
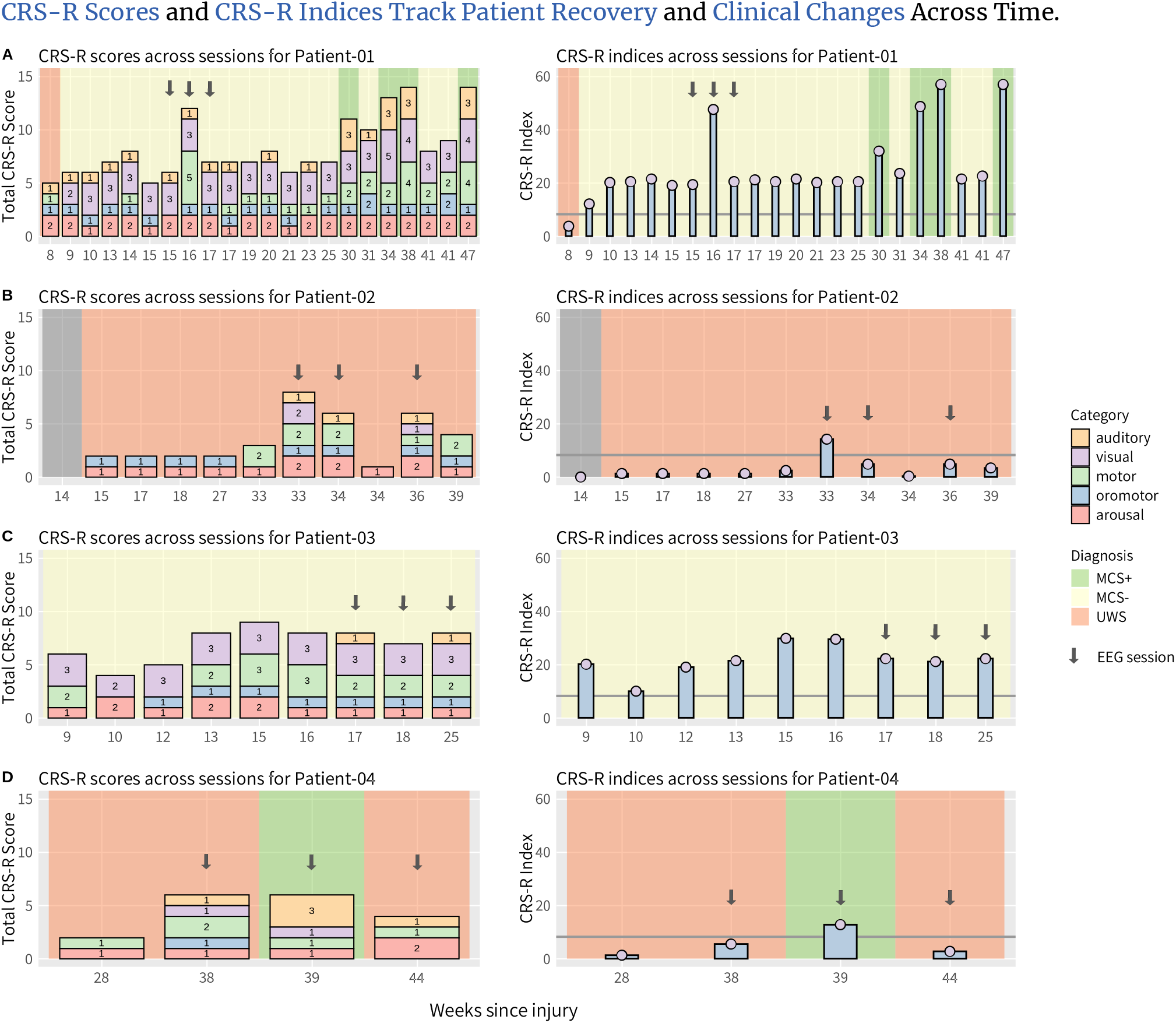
CRS-R scores and CRS-R indices across CRS-R sessions. The left column displays the total CRS-R scores, broken down into their respective sub-scores. The right column shows the CRS-R indices. Panel (A) shows the CRS-R results for Patient-01, (B) for Patient-02, (C) for Patient-03, and (D) for Patient-04. The arrows indicate the sessions during which EEG data were collected. Abbreviations: EEG: Electroencephalography; CRS-R: Coma Recovery Scale-Revised; UWS: Unresponsive Wakefulness Syndrome; MCS: Minimally Conscious state

#### Patient-01

During a period of 47 weeks post-injury, 22 sessions of the Coma Recovery Scale-Revised (CRS-R) were held, as shown in Figure 2A. The patient often demonstrated elevated arousal levels and spontaneous eye opening without external stimulation. The patient exhibited reflexive behaviors that included auditory startle responses, oral reflexive movements, and abnormal flexion/extension responses to painful stimuli. During our tests, the patient demonstrated the ability to visually track his own reflection in a mirror, which supported a diagnosis of MCS-. The CRS-R sessions conducted after our last test session show that the patient progressed significantly, transitioning to MCS+. This improvement included consistent command-following, recognizing objects, localizing objects and reaching out to them, smiling at familiar people, and being able to hold an object. However, for the purposes of our analysis, the patient was classified as MCS-, as this was the most accurate diagnosis at the time of the final EEG test.

#### Patient-02

Eleven CRS-R sessions were completed over 39 weeks post-injury (Figure 2B). Although minor improvements in the CRS-R subscores were observed during this period, the patient remained in UWS. This patient was the least responsive among all tested individuals. No eye contact was observed and the behaviors were limited to reflexive responses, such as occasional oral reflexes and pain reflexes, although these were weak. Arousal levels fluctuated between sessions; in some cases spontaneous arousal was observed without stimulation, while in others external stimulation was required.

#### Patient-03

Nine CRS-R sessions were conducted over 25 weeks post-injury (Figure 2C). The patient was classified as MCS-based on her ability to visually track herself in a mirror. The left eye did not respond due to paralysis of the eye muscles, while brief and inconsistent eye contact with the right eye was observed. Motor function was severely affected due to quadriplegia, resulting in a loss of voluntary motor control; however, reflexive responses to pain and occasional oral reflexes were preserved. Arousal levels varied between sessions; eyes opened spontaneously in some cases but required external stimulation in others.

#### Patient-04

Prior to being transferred to the collaborating hospital for testing, the patient had been diagnosed as MCS at another medical center, showing no active movements or signs of cooperation beyond reflexive responses. Four CRS-R sessions were conducted over 44 weeks post-injury (Figure 2D), during which command-following through horizontal eye movements was observed in one session, leading to a classification of MCS+. Despite this observation, overall CRS-R scores remained low, with the patient exhibiting little to no response on other tests. During EEG sessions, the patient struggled to maintain wakefulness and consistent levels of arousal. Additionally, the patient’s partner reported observing higher levels of activity outside of our testing periods, suggesting that our assessments may have coincided with lower arousal phases.

### 3.2. Spectral entropy but not neural tracking shows a relationship with the CRS-R index

Spectral entropy patterns remained relatively consistent within individual patients across multiple EEG sessions (see Appendix 1). The primary differences were observed between patients rather than within sessions for the same individual. Among the four patients, Patient-03 and Patient-04 exhibited the greatest variation in entropy across electrodes. Specifically, Patient-04 demonstrated a minimum entropy of 0.440 and a maximum of 0.746 across the three sessions (SD = 0.0567), while Patient-03 showed a minimum of 0.293 and a maximum of 0.565 (SD = 0.0532). In contrast, Patient-01 displayed the lowest variation, with entropy values ranging from 0.518 to 0.670 (SD = 0.0258). Similarly, Patient-02 showed relatively limited variation, with a minimum entropy of 0.378 and a maximum of 0.489 (SD = 0.0378).

#### Spearman Correlations

The relationship between spectral entropy and the CRS-R index exhibited a square root pattern, which was more pronounced during listening conditions (Fig. 4 A-B) and less evident during baseline recordings (Fig. 4 C). In the Dutch listening condition, the spectral entropy demonstrated significant Spearman correlations with the CRS-R index: 0.65 (*p* ≤ 0.001) for 5 minutes of data and 0.71 for 6 minutes (*p* ≤ 0.001) and 10 minutes (*p* ≤ 0.001) of data. Similarly, the Swedish listening condition yielded significant Spearman correlations of 0.57 (*p* ≤ 0.001) for 5 minutes of data and 0.56 (*p* ≤ 0.001) for 6 minutes of data. In contrast, the baseline condition did not show a significant Spearman correlation (*r* = 0.29, *p* = 0.132). The neural tracking of the speech envelope showed no relationship with the CRS-R index. In the Dutch listening conditions, the spearman correlation was 0.06 (*p* = 0.805) for 6 minutes of data and 0.03 (*p* = 0.912) for 10 minutes of data, while in the Swedish listening condition it was 0.04 (*p* = 0.876), see Figure 3.

**Figure 3.**
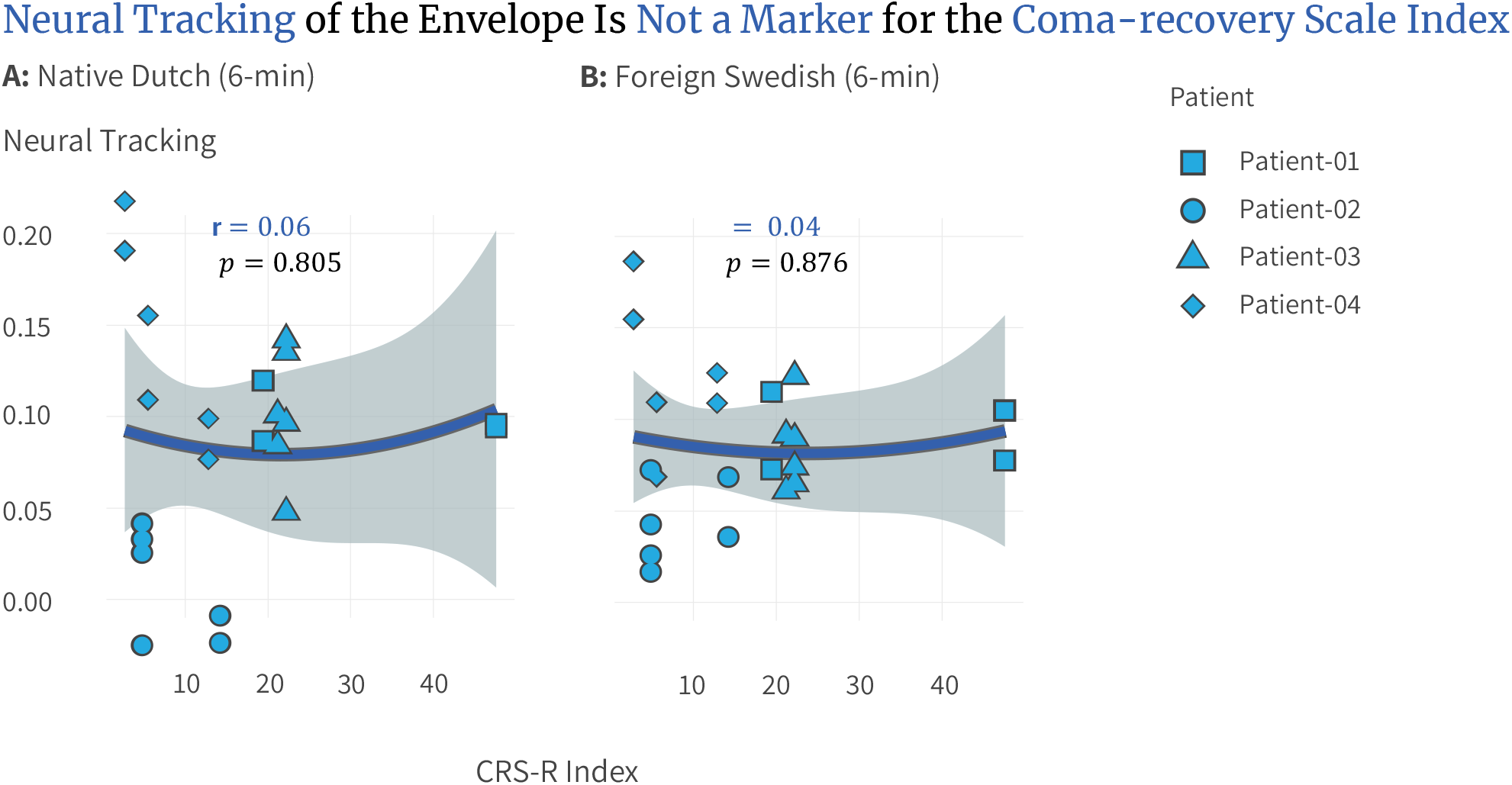
Relationship between neural tracking and the CRS-R index. Each dot represents one session. (A) For Dutch listening data, both 6-minute and 10-minute segments exhibit similar patterns, with Spearman correlations of 0.06 (6-min) and 0.03 (10-min). (B) For Swedish listening data, the 6-minute segment shows a Spearman correlation of 0.04. All correlations are non-significant. The smooth line represents a quadratic (second-degree) fit, included solely for visualization purposes to highlight potential non-linear trends in the data; Abbreviation: CRS-R: Coma Recovery Scale-Revised.

**Figure 4.**
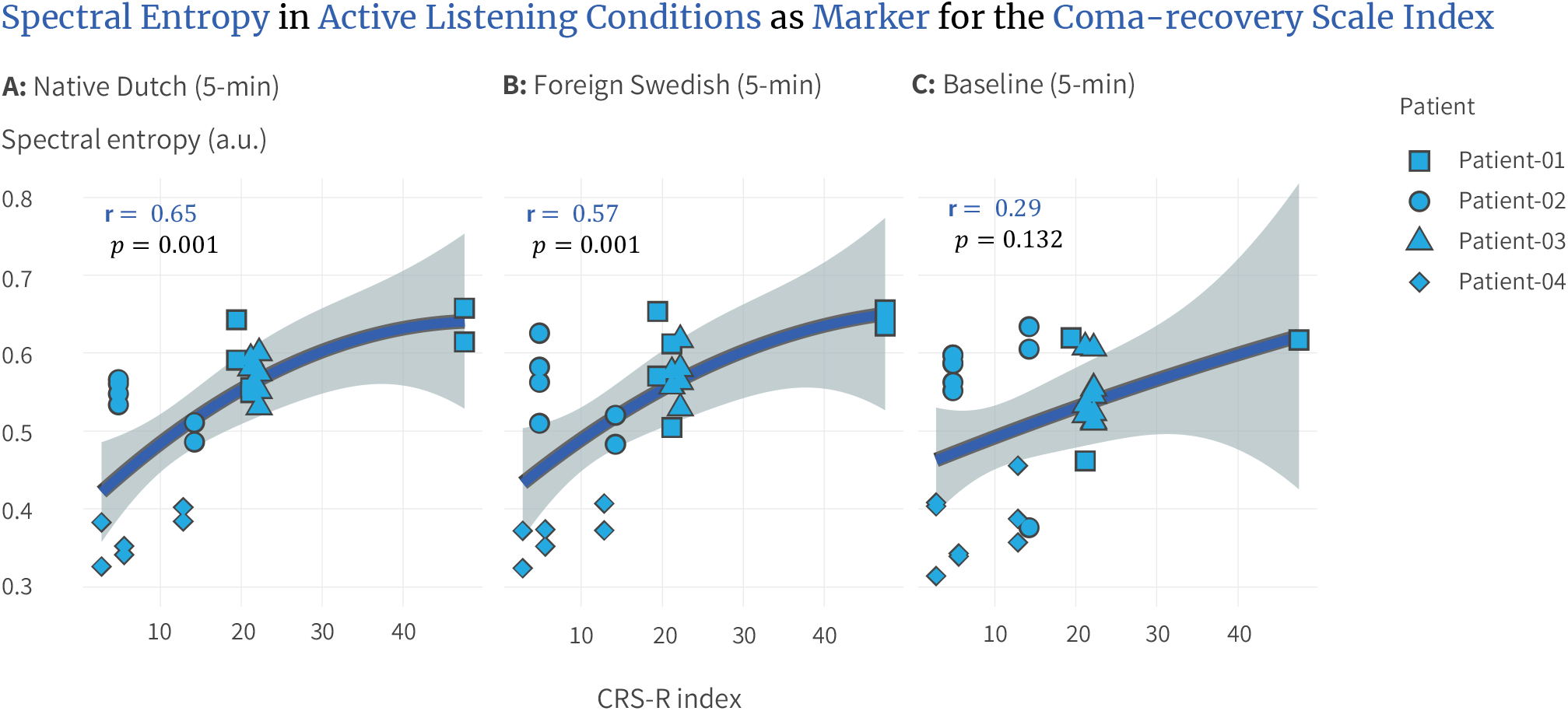
Relationship between spectral entropy and the CRS-R index. Each dot represents one session. The Dutch and Swedish conditions were trimmed to match the baseline duration. (A) For Dutch listening data, the 5-minute, 6-minute, and 10-minute segments show similar patterns, with significant Spearman correlations of 0.65 (5-min), 0.71 (6-min), and 0.71 (10-min). (B) For Swedish listening data, the 5-minute and 6-minute segments also exhibit similar patterns, with significant Spearman correlations of 0.57 (5-min) and 0.56 (6-min). (C) The baseline condition shows a non-significant Spearman correlation of 0.29. The smooth line represents a quadratic (second-degree) fit, included solely for visualization purposes to illustrate potential non-linear trends in the data. Abbreviation: CRS-R: Coma Recovery Scale-Revised, au.: arbitrary units.

#### BRM Results

The individual predictor models— spectral entropy (Eq. 4) and neural tracking (Eq. 5)—produced posterior estimates with 95% credible intervals (CI) that included zero, indicating insufficient evidence to confidently establish their independent effects (Table 3). For spectral entropy, 94.33% of the posterior distribution was positive, with a mean estimate of *β*_*SpE*_ = 5.5, suggesting a potential positive association supported by the correlation plots. In contrast, neural tracking showed 76.55% of its posterior distribution as negative, with a smaller mean estimate of *β*_*NT*_ = − 3.34. Combined with the lack of correlation between neural tracking and the CRS-R index, this points to the likelihood that neural tracking does not have a meaningful effect in this context.

**Table 3:**
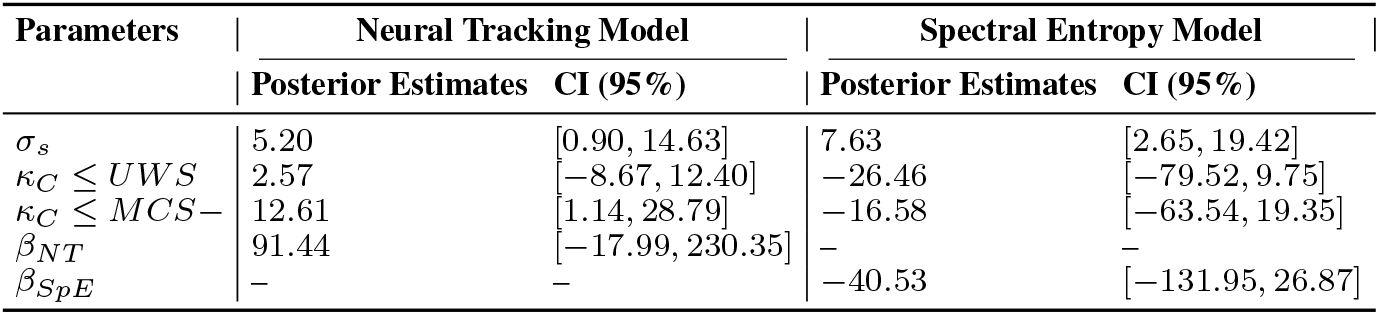
Results from the spectral entropy model (Eq. 7) and the neural tracking model (Eq. 8). The table shows the posterior estimates and 95% credible intervals(CI) for each parameter across the two models.

The interaction model (Eq. 6) provided evidence of an interaction effect between neural tracking and spectral entropy, with a substantial posterior estimate of 89.47 (95% CI: [10.03, 171.39]). Furthermore, the model revealed evidence of a negative main effect of neural tracking (*β*_*NT*_ = − 40.72, 95% CI: [− 82.37, − 0.91]). To improve the interpretability of this interaction and since both predictors are continuous, we visualized their relationship using anchor points (Figure 5). The visualization illustrates that higher neural tracking values amplify the positive relationship between spectral entropy and CRS-R indices. In other words, patients with stronger neural tracking exhibit a more pronounced positive association between spectral entropy and CRS-R indices.

**Figure 5.**
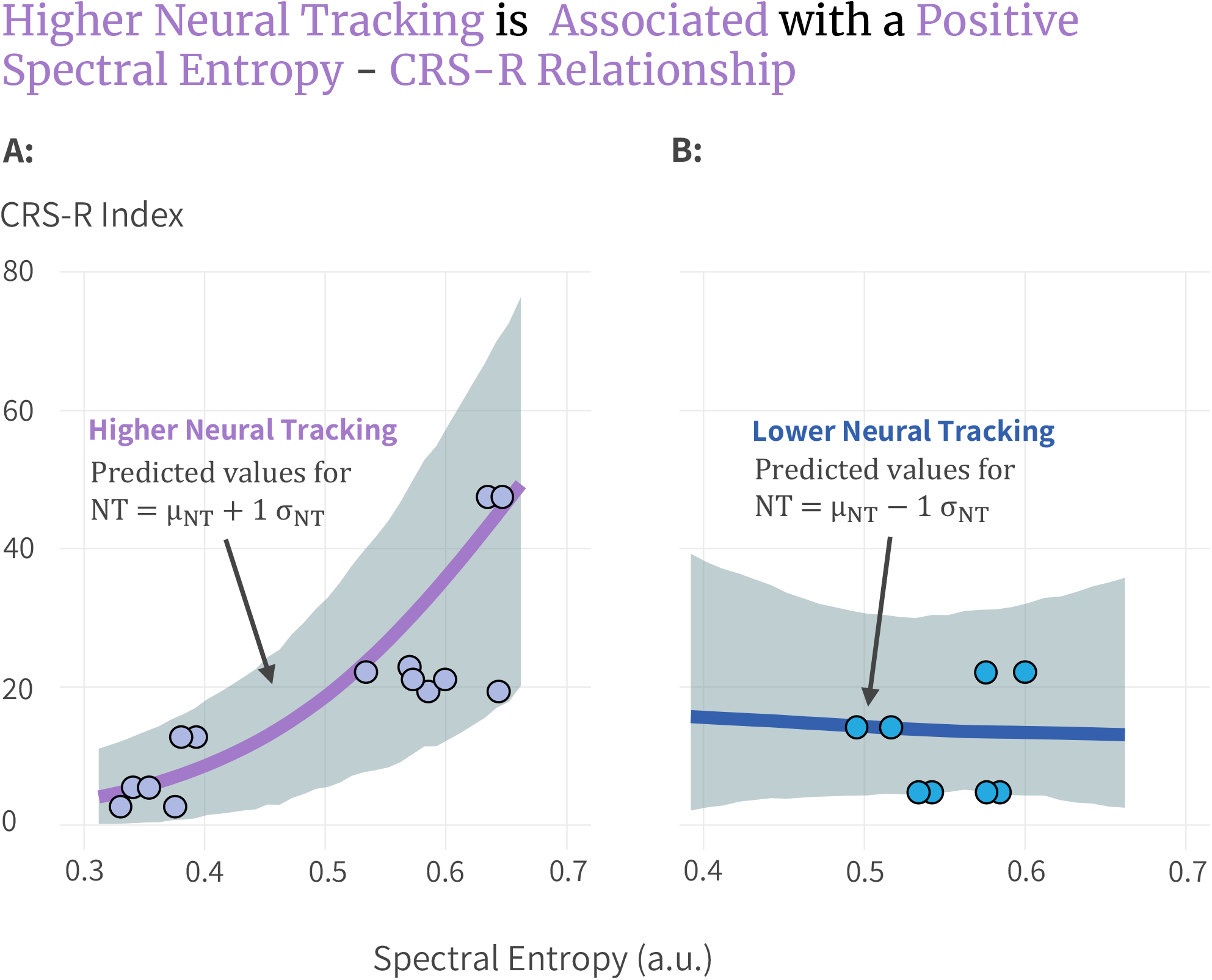
The interaction effect between neural tracking (NT) and spectral entropy on the CRS-R index using a Bayesian regression model. Since both spectral entropy and neural tracking are continuous variables, two representative NT values were selected to visualize the interaction effect: Higher neural tracking corresponds to the mean NT value plus one standard deviation (NT = μ_NT_ + 1s_NT_). Lower neural tracking corresponds to the mean NT value minus one standard deviation (NT = μ_NT_ −1s_NT_).(A) For higher neural tracking, a positive relationship is observed between spectral entropy and the CRS-R index, as shown by the curved Bayesian regression line (purple) and its 95% credible interval (shaded area). Each dot represents real patient session data. While individual NT values vary continuously, they are anchored to either high or low NT conditions for visualization purposes. The smooth lines highlight the modeled interaction effect between spectral entropy and NT on CRS-R index. Abbreviations: CRS-R: Coma Recovery Scale-Revised; NT: Neural Tracking; μ_NT_ : Mean Neural Tracking; s_NT_ : Standard Deviation of Neural Tracking, a.u.: arbitrary units.

### 3.3. Neural tracking shows a potential relationship with the CRS-R diagnosis

#### Boxplots

Figure 6 illustrates the distribution of neural tracking and spectral entropy values in diagnostic categories (UWS, MCS-, and MCS+). Neural tracking shows minimal overlap between groups, with a clear trend of increasing values as diagnostic severity decreases. In contrast, while the spectral entropy also displays minimal overlap between categories, it does not follow the expected pattern of increasing values with decreasing severity. In particular, the MCS+ category exhibits the lowest spectral entropy values; however, it is important to consider that this result is driven by a single MCS+ patient, which may limit the generalizability of this observation.

**Figure 6.**
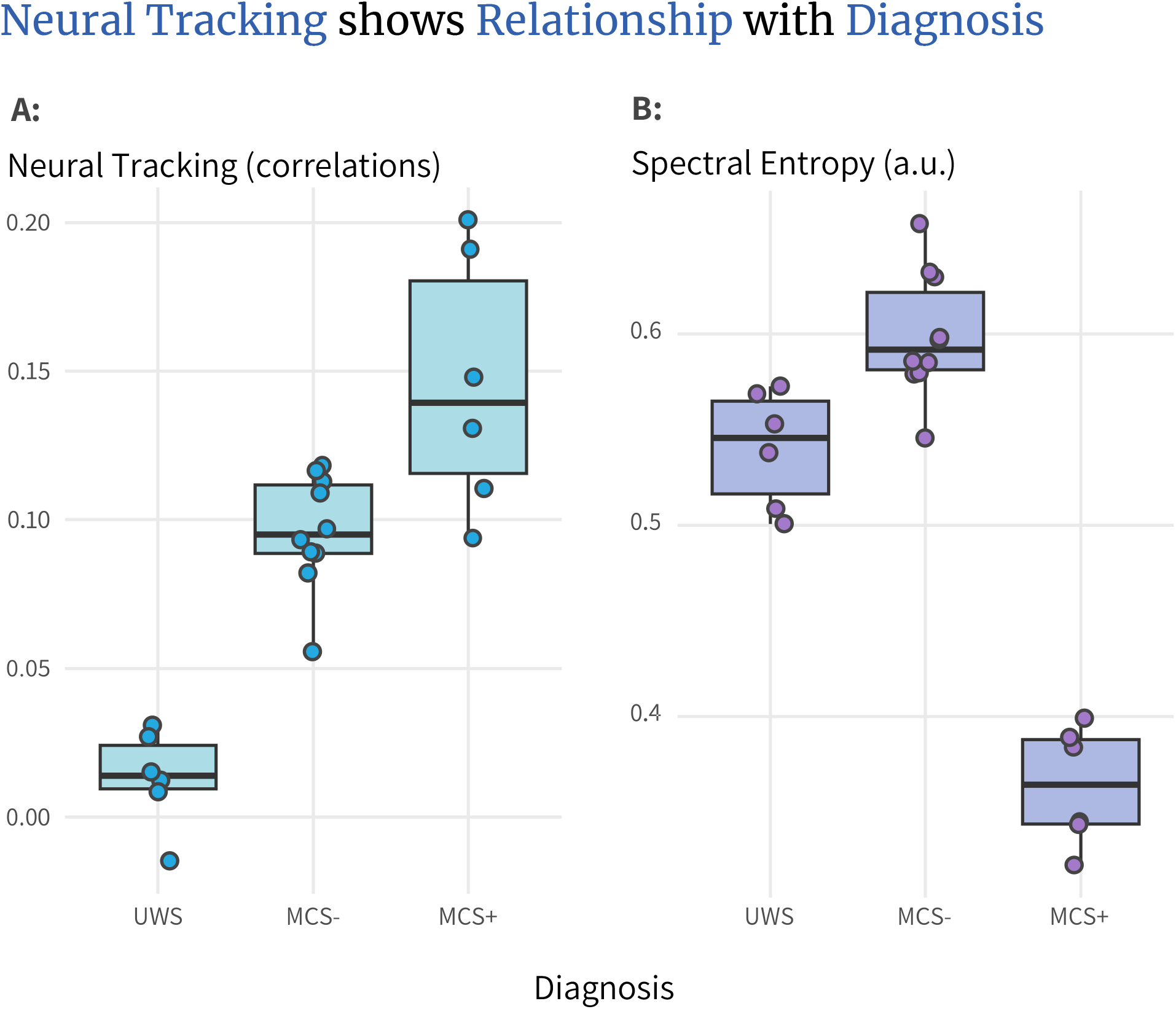
Patients are grouped according to their best achieved diagnostic classification across three assessment sessions. Each dot represents the neural responses from a single listening condition.(A) shows the neural tracking values distributed across diagnostic categories (B) shows the spectral entropy distributed across diagnostic categories. Abbreviations: UWS: Unresponsive Wakefulness Syndrome; MCS: Minimally Conscious state, a.u.: arbitrary units

#### BRM results

The linear relationship observed in Figure 6A is quantitatively captured by our Bayesian regression models. The neural tracking model (Eq. 8) showed a substantial posterior estimate (*β*_*NT*_ = 91.44, 95% CI −17.99, 230.35), with 95.72% of the posterior distribution being positive. This indicates a likely positive relationship between neural tracking and diagnosis, although the wide CI reflects considerable uncertainty in parameter estimation. Specifically, there was evidence supporting the threshold parameter (*κ*_*C*_ ≤ MCS-), which distinguishes UWS and MCS-from MCS+ (*κ*_*C*_ = 12.61, 95% CI: 1.14, 28.79). In contrast, the threshold (*κ*_*C*_ ≤ UWS) is unable to distinguish UWS from both MCS- and MCS+(*κ*_*C*_ = 2.57, 95% CI: −8.67, 12.40).

Using posterior sampling of the parameter distributions, the relationship between neural tracking, spectral entropy, and diagnosis is visualized in Appendix 2. The analysis indicates that, for this sample, the 50% probability threshold for transitioning from UWS to MCS- occurs at approximately 0.04 NT, while the transition from MCS-to MCS+ is observed around 0.13 NT. For neural tracking values that exceed 0.10, the model demonstrates greater certainty that higher NT values are unlikely to be associated with UWS, as reflected by narrower CIs. Similarly, for MCS+, the model shows greater certainty in the opposite direction. However, for MCS-, uncertainty remains relatively high across neural tracking values, suggesting less precision in distinguishing this category. In contrast, spectral entropy exhibits consistently large CIs across all diagnostic categories, reflecting greater variability and reduced certainty in its predictive capacity. This is likely influenced by the single MCS+ patient, whose data disrupt the expected linear trend of spectral entropy values.

## 4. Discussion

Our analysis revealed evidence of a relationship between spectral entropy and the CRS-R index, supporting an interaction effect between spectral entropy and neural tracking. Specifically, patients with higher neural tracking showed a stronger association between spectral entropy and the CRS-R index. The results further indicate that the association between spectral entropy and the CRS-R index is stronger when patients listen to a story, in their native language or a foreign language, compared to a baseline condition.

The neural tracking itself did not show evidence of a direct relationship with the CRS-R index. However, there was evidence of its association with the diagnostic categories of the CRS-R. Patients who achieved a diagnosis of MCS during at least one CRS-R session displayed higher neural tracking values. Patient-04, diagnosed as MCS+, exhibited the highest neural tracking value, while Patient-01 and Patient-03, diagnosed as MCS-, showed intermediate to strong values, and Patient-02, diagnosed as UWS, showed the lowest neural tracking.

It is important to note that this study involved a small sample of four patients, although with multiple sessions per patient. In Bayesian statistics, small sample sizes amplify the influence of priors on posterior distributions. We used non-informative priors to allow the models greater flexibility, which led to wider CIs. In Bayesian statistics, CIs that include zero do not necessarily indicate the absence of an effect; they reflect both the use of uninformative priors and genuine uncertainty about the magnitude of the effect. Future studies with larger samples are needed to confirm these findings and provide greater certainty.

### 4.1. interaction effect

The observed interaction between neural tracking and spectral entropy is intriguing. It suggests that patients with stronger neural tracking may have auditory cortices better adapted to processing complex signals, leading to entropy values that could more precisely reflect patients’ behavioral state. However, drawing definitive conclusions about the underlying mechanisms remains premature given the limitations of the current sample size.

### 4.2. Spectral entropy

Patients in this study showed SpE values consistent with those reported in previous research. For direct comparison, it is necessary to transform the SpE values, as the Entropy™ algorithm (Viertiö-Oja et al., 2004) applies a nonlinear spline function to convert entropy values from 0–1 to a 0–91 scale. In our dataset, the mean SpE across all electrodes ranged from 0.293 to 0.612, corresponding to approximately 15–50 on the Entropy™ scale. Some individual electrodes reached values as high as Gosseries et al., 2011 reported that, among chronic patients with DoC (1 *>* month post-injury), those with UWS had mean Entropy™ values of 45±28, with the lowest scores around 10. Patients with MCS had mean values of 73±19, with the lowest around 30. In healthy controls, Entropy™ values typically ranged from 80 to 89 (Gosseries et al., 2011; Vakkuri et al., 2004).

Studies have shown that entropy measures correlate with CRS-R scores (Gosseries et al., 2011; Visani et al., 2022). In this study, we demonstrate that there is likely a similar relationship between spectral entropy and the CRS-R index, a recalculated version of the CRS-R score designed to provide a more interpretable and diagnostically meaningful measure (Annen et al., 2019). An open question arises with Patient-04, who demonstrated command-following (diagnosed as MCS+) but consistently exhibited very low spectral entropy compared to other patients. At the group level, higher entropy values are generally more prevalent in patients with MCS compared to those with UWS. However, this trend does not necessarily hold at the individual level. Significant variability in entropy values remains among patients with MCS. In particular, Patient-04’s highest entropy value (0.565) exceeded the maximum entropy recorded for Patient-02 (diagnosed with UWS), whose maximum was 0.489. This discrepancy may reflect testing during a low arousal phase, supported by clinical observations such as repeated awakening protocols and reports from the patient’s partner on reduced responsiveness on test days. Previous research indicates that patients with MCS experience fluctuations between high and low arousal states (Wannez et al., 2017). These fluctuations may be related to the variability of the spectral entropy (Piarulli et al., 2016).

### 4.3. Neural tracking

In addition to spectral entropy, we have demonstrated that using a backward model for the neural tracking of the speech envelope can be used as an effective tool to investigate speech processing in patients with DoC. Although neural tracking is known to diminish when attention to stimuli is reduced, it remains detectable under such conditions (Dieudonné et al., 2025; Geirnaert et al., 2021; Vanthornhout et al., 2019). Our findings further reinforce that neural tracking can be reliably measured in patients with DoC, and for patients with MCS achieving neural tracking values comparable to those observed in healthy individuals in EEG (e.g. Vanthornhout et al., 2018) and MEG (e.g., Ding & Simon, 2012), even when they appear drowsy and have difficulty staying awake.

Although previous research has shown that patients with MCS-generally exhibit reduced connectivity between Broca’s region and other language-related cortices (Bruno et al., 2012), and are behaviorally characterized as lacking signs of language processing. Our findings suggest that, despite their severe impairments, MCS-patients may retain some capacity to process the sensory-driven automatic aspects of language input. This aligns with the work of Jia et al., 2023, who proposed that the envelope-tracking response to speech can be generated by automatic processes that are minimally influenced by the state of consciousness. Our results further support this interpretation: spectral entropy patterns were similar for both Dutch (the native language) and Swedish (the foreign language), and we found neural tracking for both Dutch and Swedish, indicating that certain neural responses are preserved regardless of language comprehension. However, while neural tracking can be observed even in response to incomprehensible speech, the presence of comprehension can enhance the strength of neural tracking responses (Chen et al., 2023; Gillis et al., 2023; Kösem et al., 2023). Several other studies have also found evidence of residual language processing in patients with DoC. Patients with MCS have been shown to exhibit a higher phase-locking index and phase coherence to sound rythms compared to those with UWS, with these measures showing a positive correlation with CRS-R scores (Binder et al., 2017, 2020; Górska & Binder, 2019). Furthermore, phase coherence with both rythmic sounds and natural speech has been found to be higher in patients who demonstrate clinical improvements over a 6-month period, also showing a positive correlation with CRS-R scores (Xu et al., 2021). Gui et al., 2020 further investigated linguistic processing in patients with DoC by assessing their ability to track linguistic structures such as words, phrases, and sentences. Their findings revealed that patients with MCS can retain some degree of higher-level linguistic processing. A key distinction between our study and previous research is that we differentiated between MCS+ and MCS-groups, while none of the aforementioned studies made this distinction.

Neural tracking using a decoder model offers two key advantages over previously explored auditory methods in DoC: its versatility in using various speech features such as the speech envelope and the fundamental frequency of the voice (*f* 0) for the decoder model (Van Canneyt et al., 2021) and linguistic features based on phoneme and words for the encoder model (e.g., Di Liberto et al., 2015; Gillis et al., 2021, 2022; Puffay, Vanthornhout, et al., 2023). Among these features, the speech envelope is of particular importance because it aligns closely with how the brain perceives and interprets speech signals after processing by the ear and cochlea (Ding et al., 2014), and plays an important role in speech intelligibility (Shannon et al., 1995; Vanthornhout et al., 2018). Using a decoder model, we can establish a direct mapping between neural activity and the speech envelope, enabling us to quantify how effectively the brain “tracks” this envelope. Thus, neural tracking offers an ecologically relevant measure of speech processing in patients with DoC. These capabilities make neural tracking a powerful tool for assessing residual language abilities in patients with DoC. As demonstrated by its successful application in other domains, such as aphasia research (De Clercq et al., 2025; Kries et al., 2024; Mehraram et al., 2024)

In addition to its ecological validity, we have demonstrated the feasibility of using a mobile EEG system that can be brought directly to the patient’s bedside. This portability offers convenience by eliminating the need for patients to travel to specialized rooms or facilities. Moreover, presenting naturalistic stories can provide a more engaging and pleasant experience for patients. Together, these advantages make this approach a practical and patient-friendly tool to evaluate residual language processing in clinical settings.

### 4.4. Residual language assessment

Research on language assessment and finding residual language processing in patients with DoC is gaining momentum (for a review see Aubinet et al., 2022). Not only can detecting residual language processing help with improving the diagnosis, but it also raises awareness that non-communicating patients may still process language to some degree. Patients who appear to be unconscious may still have internal experiences that cannot be expressed outward (Lawrence et al., 2023). In fact, in one study, 27% of patients who were unconscious during hospitalization reported being able to hear and understand their surroundings at times. (Lawrence, 1995). In another study, 8 out of 15 patients with posttraumatic coma had some degree of memory of being comatose (Tosch, 1988). Patients who do not respond typically receive less verbal communication and interaction compared to those who are verbally responsive (Alasad & Ahmad, 2005; Happ, 2021), leading to dehumanization of these patients (Lekka et al., 2021). These studies underscore the clinical importance of auditory and language paradigms in DoC. These approaches not only have the potential to improve diagnostic accuracy but also emphasize the need to engage verbally with all patients, regardless of their responsiveness, acknowledging that they may have a hidden internal life.

### 4.5. Directions for future research

This study provides initial evidence that neural tracking of the speech envelope can be measured in patients with disorders of consciousness. However, many questions remain unanswered, which paves the way for future research. Replicating these findings with larger and more diverse patient data sets will be essential to validate and generalize the results. In addition, other speech features could be modeled using the decoder and encoder approach or deep neural networks (Puffay, Accou, et al., 2023). Investigating these features may offer deeper insight into the cognitive and linguistic processing of patients. Exploring longer durations of story listening (for example, *>* 1 hour) could also yield valuable information, as patients with MCS are known to exhibit cycles of arousal. Understanding how neural tracking fluctuates across these arousal cycles and compares with spectral entropy would provide a more dynamic view of the neural responses of patients. Although this study focuses mainly on diagnostic applications, the prognostic value of neural tracking is not yet explored. Future research should investigate whether neural tracking can predict recovery trajectories or long-term outcomes in patients with disorders of consciousness. These avenues of research hold promise for advancing both our understanding and clinical applications of neural tracking in this population.

## 5. Conclusion

This study provides evidence of a relationship between spectral entropy and the CRS-R index, with higher neural tracking associated with a stronger link, particularly during story listening conditions. Neural tracking was also associated with the diagnostic categories of CRS-R, which increased with better diagnostic outcomes. Although these findings are encouraging, they are based on a small sample size, highlighting the need for future research with larger datasets to validate and expand upon these results.

## 6. Data availability

The data underlying this article cannot be shared publicly due to privacy and ethical restrictions related to the protection of individuals who participated in the study. Access is limited to authorized collaborators and regulatory authorities under strict confidentiality, as outlined in the informed consent. Data may be shared upon reasonable request to the corresponding author and subject to institutional and ethical approval.

## 8. Acknowledgements

This study is funded by the Flemish scientific research fund (FWO; G0D6720N and 1290821N). The authors thank Aurore Thibaut, Charlène Aubinet, Emilie Szymkowicz, and Marie Vitello of the Coma Science Group for their guidance and support in teaching us the CRS-R assessment. The authors also extend thanks to Erika Van Den Steen for narrating the story in both Dutch and Swedish, and to Anke Van Dijck from AZ Klina for facilitating the research collaboration.

## 10. Appendix A: Bayesian Regression Model Specifications

### CRS-R diagnosis model

Once the model is fitted, the regression coefficients can be used to predict the cumulative probabilities for each ordinal category using the inverse logit function. Subsequently, the posterior probability of a neural tracking or spectral entropy value belonging to a specific diagnosis category can be expressed as:

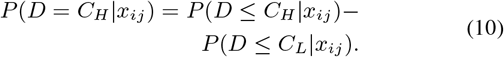

This expression can be computed for a specific subject or marginalized across subjects, ignoring random subject effects.

where:

- *x*_*ij*_ is the *i*-th measurement value, such as neural tracking or spectral entropy, for individual *j*.
- *D* represents the diagnosis.
- *P* (*D* = *C*_*H*_| *x*_*ij*_) is the posterior probability that the diagnosis *D* corresponds to category *C*_*H*_, given the observed measurement *x*_*ij*_.
- *P* (*D*≤ *C*_*H*_ | *x*_*ij*_) is the cumulative posterior probability that the diagnosis *D* falls into category *C*_*H*_ or any lower ordinal category, given measurement *x*_*ij*_.
- *P* (*D* ≤ *C*_*L*_| *x*_*ij*_) is the cumulative posterior probability that the diagnosis *D* falls into category *C*_*L*_, which is ordinarily ranked lower than *C*_*H*_, given the measurement *x*_*ij*_.

### Priors

In each model, we used the default priors specified by the ‘brms’ package, see Table 4. These priors impose minimal constraints while allowing the data to guide the parameter estimation.

**Table 4:** Default priors chosen by the brms package.

| Parameter | Prior Distribution |
| --- | --- |
| $\beta$ | Flat($-\infty, +\infty$) |
| $s_j$ | $N(0, \sigma_u^2)$ |
| $\sigma_s$ | $t(3, 0, 2.5)$ |
| $\kappa_C$ | $t(3, 0, 2.5)$ |
| $\phi$ | Gamma(0.01, 0.01) |

### Model diagnostics

All Bayesian regression models were estimated using Markov Chain Monte Carlo (MCMC) sampling with 4 parallel chains, each running 10,000 iterations (including 5,000 warmup iterations). Convergence was assessed through multiple diagnostics: all models except model 9 achieved the recommended 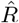 values of ≤ 1.01, indicating successful chain convergence. The successful models maintained effective sample sizes exceeding 400 for all parameters, ensuring reliable posterior estimation. Visual inspection of traceplots confirmed consistent mixing patterns across all chains.

**Figure Appendix 1:**
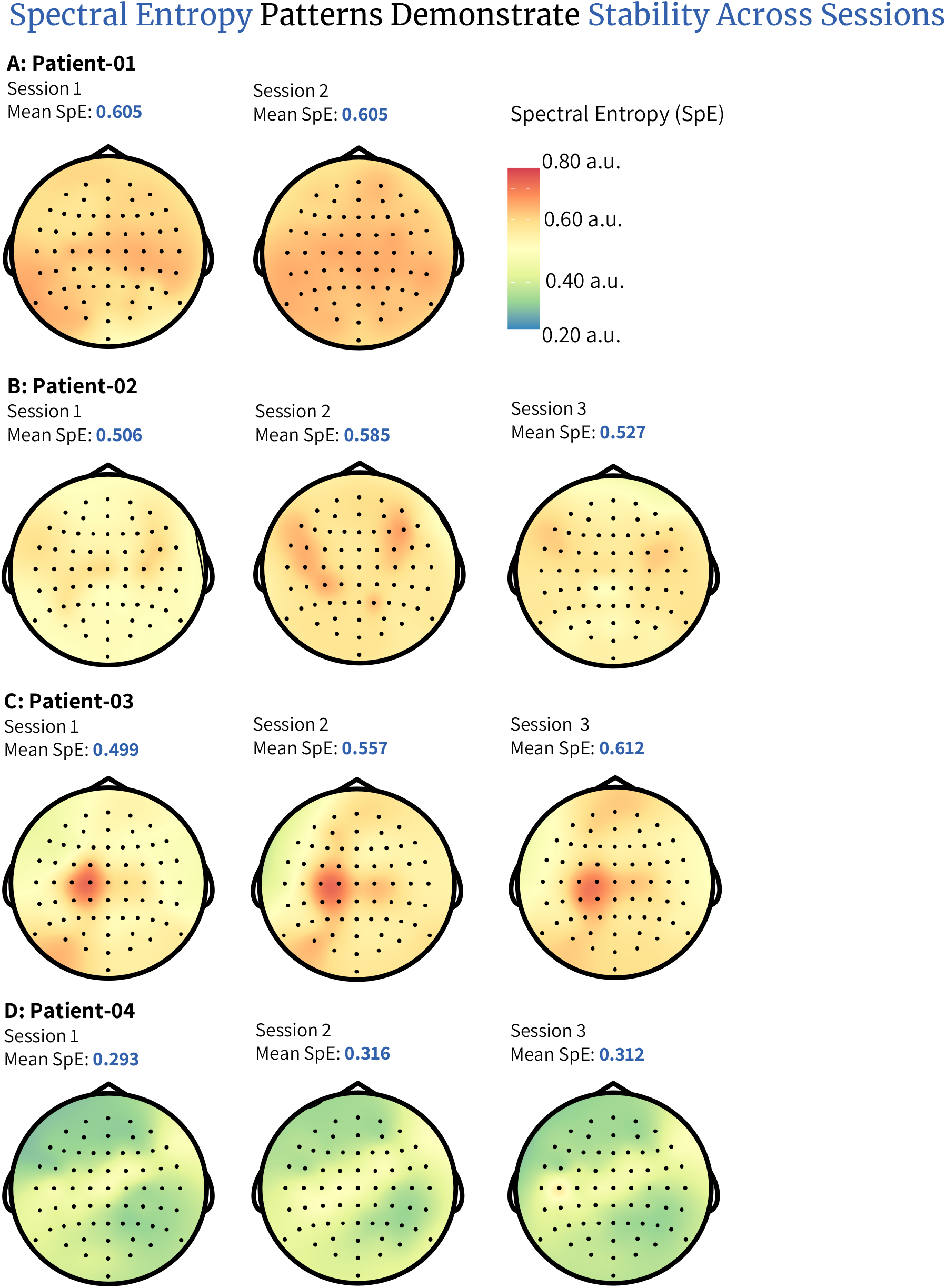
Spectral entropy averaged per session across conditions for each patient. Panel (A) shows the spectral entropy for Patient-01, panel (B) for Patient-02, and panel (C) for Patient-03 (D) for Patient-04; Abbreviations: SpE: Spectral Entropy, a.u.: arbitrary units

**Figure Appendix 2:**
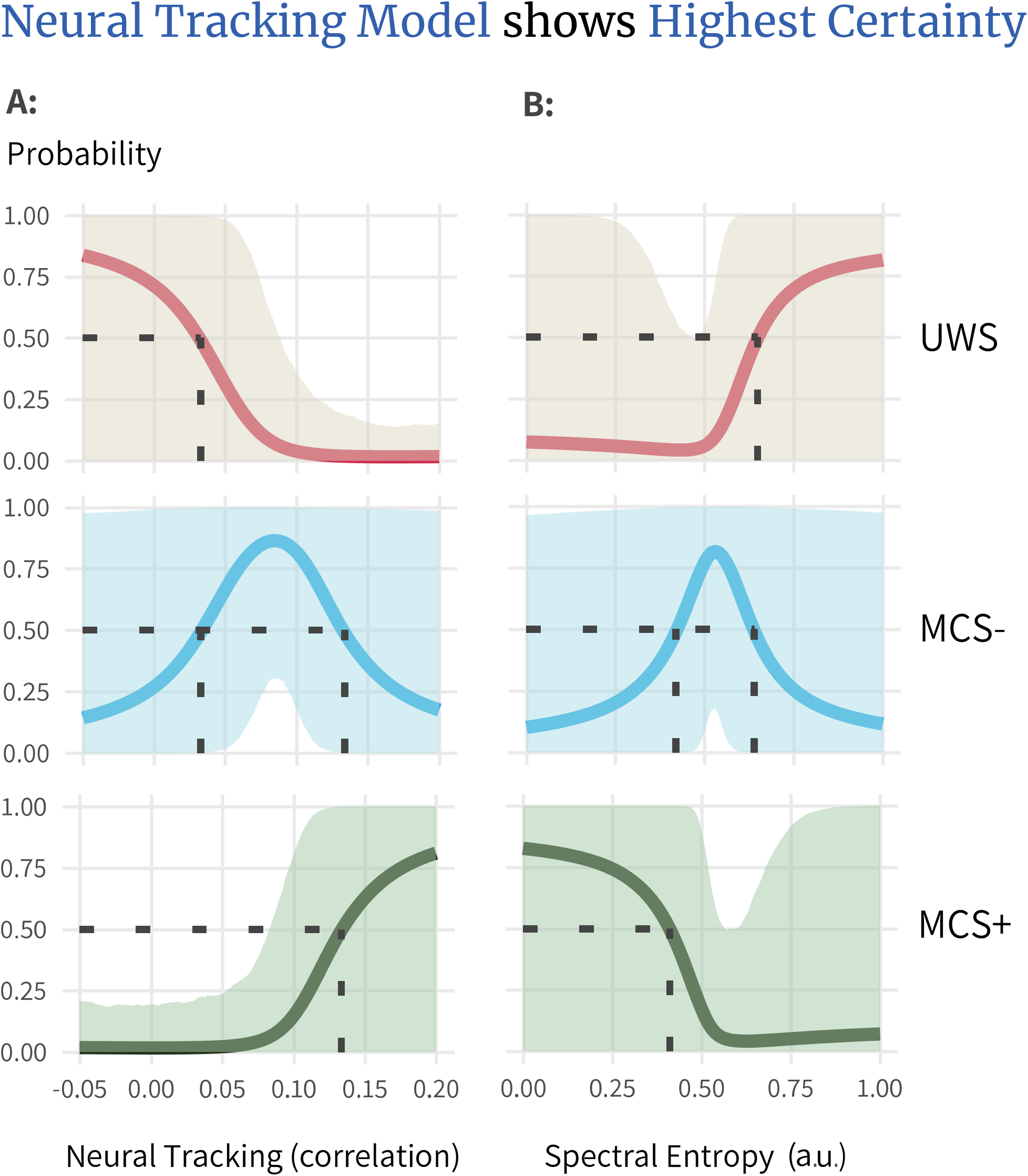
This figure shows the estimated predicted probabilities of each diagnostic category based on (A) neural tracking and (B) spectral entropy values using ordinal Bayesian models 7 and 8. The predictions are population-level rather than subject-specific. The probabilities are shown by their mean (solid line), as well as their 95% credible intervals (shaded area). Abbreviations: UWS: Unresponsive Wakefulness Syndrome; MCS: Minimally, a.u.: arbitrary units

